# REDUCED ALPHA-BAND PHASE COHERENCE AND CORTICAL COMPLEXITY IN FIBROMYALGIA: A TMS-EEG EXPLORATORY STUDY

**DOI:** 10.1101/2025.09.29.678190

**Authors:** Anne Jakobsen, Enrico De Martino, Bruno A. N. Couto, Margit M. Bach, Stian Ingemann-Molden, Thorvaldur S. Palsson, Jakob Udby Blicher, Adenauer G. Casali, Thomas Graven-Nielsen, Daniel Ciampi de Andrade

**Author notes:** **Corresponding author** Daniel Ciampi de Andrade Center for Neuroplasticity and Pain (CNAP), Department of Health Science and Technology. Faculty of Medicine, Aalborg University Selma Lagerløfs Vej 249 room 12.02.018. Gistrup 9260 Denmark.

## Abstract

**Objectives:** Cortico-spinal excitability of the primary motor cortex (M1) is reduced in fibromyalgia, and repetitive transcranial magnetic stimulation (TMS) targeting M1 normalizes these changes and relieves symptoms. TMS combined with electroencephalography (TMS-EEG) allows the measurement of M1 excitability and its connectivity to other regions, which may help clarify neurophysiological effects in fibromyalgia. We assessed cortical excitability, oscillatory activity, and complexity in individuals with fibromyalgia compared to pain-free healthy controls.

**Methods:** Global and local mean field power, peak-to-peak amplitude, event-related spectral perturbation, intertrial coherence (ITC), natural frequency, and perturbational complexity index (PCIst) of the EEG response after left-M1 TMS were compared between groups (n=18 fibromyalgia; n=15 controls). Pain intensity, interference, relief of current therapy, mood, and quality of life were assessed in individuals with fibromyalgia.

**Results:** Compared with controls, individuals with fibromyalgia showed a reduction in the alpha-band ITC in middle and right parieto-occipital areas (P<0.05). Middle-parieto-occipital ITC negatively correlated with reported pain relief (rho=-0.552, p=0.019). The PCIst was lower in fibromyalgia compared with controls (P<0.01) and correlated with higher pain interference in general activity (rho=-0.486, p=0.042).

**Conclusion:** Individuals with fibromyalgia showed abnormal cortical connectivity compared with asymptomatic controls.

**Significance:** TMS-EEG measurements may provide insights on brain connectivity relevant for therapy.

## 1. INTRODUCTION

Fibromyalgia (FM) is a primary pain disorder characterized by generalized widespread pain (Wolfe et al., 2016), affecting approximately 2-4 % of the general population (Heidari et al., 2017). The FM etiology and pathophysiology are poorly understood, and both peripheral and central mechanisms have been proposed (Jurado-Priego et al., 2024). However, peripheral abnormalities observed in individuals with FM do not correlate with symptoms (Kaziyama et al., 2020; Marshall et al., 2024). In contrast, stronger evidence suggests that abnormal central nervous system processing of nociceptive information contributes to FM (Jurado-Priego et al., 2024). For instance, structural brain alterations, such as decreased grey matter volume in the cingulate and prefrontal cortices (Shi et al., 2016), has been described in individuals with FM. Importantly, FM exhibits altered primary motor cortex (M1) corticospinal excitability, such as reduced intracortical inhibition (Kaziyama et al., 2020; Mhalla et al., 2010), which correlate with clinical characteristics, such as fatigue, catastrophizing, and depression (Mhalla et al., 2010) and which can partially be restored by therapy with repetitive transcranial magnetic stimulation (rTMS) to M1 (Mhalla et al., 2011). Together, this indicates that brain networks including M1 may be affected in FM.

In addition to the well-known motor functions, M1 is deeply connected to top-down pain modulatory systems, cognitive, and allostasis control (Gordon et al., 2023). Its modulation by epidural (Fonoff et al., 2009) and transcranial neuromodulatory approaches (de Andrade et al., 2011) are known to engage endogenous opioid receptors, release dopamine in the striatum, and to influence nociceptive processing in the thalamus and spinal cord (França et al., 2013; Pagano et al., 2012), for a review see (Ciampi de Andrade & García-Larrea, 2023; Moisset et al., 2016). Therefore, measures of brain connectivity after probing M1 may reveal functional connections and interactions between this area and other, distant cortical regions, providing new insights into the pathophysiology of FM and opening possibilities for improving clinical management (De Martino, Nascimento Couto et al., 2025). The recent coupling of TMS with simultaneous electroencephalography recording (TMS-EEG) has allowed for the measurement of cortical excitability and connectivity with high temporal resolution by directly stimulating specific cortical areas (Ilmoniemi et al., 1997). The TMS-EEG technique allows for the assessment of cortical excitability with measures such as global mean field power (GMFP), local mean field power (LMFP), and TMS-evoked potentials (TEPs) (Tremblay et al., 2019). As such, it can be used to estimate oscillatory dynamics, such as event-related spectrum perturbation (ERSP, a measure of changes in power across frequencies following a probing stimulus) and intertrial coherence (ITC, a measure of regional phase consistency across trials) (Makeig et al., 2004; Tremblay et al., 2019). During experimental pain in pain-free controls, reduced GMFP, LMFP, and peak-to-peak amplitude have been described (De Martino et al., 2023), as well as an ITC reduction in the alpha (8-12 Hz) frequency band (De Martino et al., 2024). Similar reductions in oscillatory frequency (as measured by the natural frequency of the oscillations) have been observed in clinical disorders such as depression, bipolar disorder, and schizophrenia (Canali et al., 2015), with a recent study reporting that these changes are associated with severity of symptoms (Donati et al., 2025). Additionally, TMS-EEG enables assessment of the complexity of evoked TMS pulses, using metrics, such as the perturbational complexity index based on state transitions (PCIst). PCIst is a metric that has been both theoretically grounded and empirically validated as an index of the capacity to sustain conscious experience in both healthy individuals and patients recovering from coma(Casali et al., 2013; Casarotto et al., 2016; Comolatti et al., 2019).

In this explorative study, TMS-EEG was used to assess cortical excitability, TMS-evoked oscillatory activity, and PCIst in individuals with FM compared to pain-free healthy controls. The overarching aim of this exploratory study was to generate hypotheses that could later be tested in further studies. First, we investigated whether decreases in cortico-spinal excitability assessed by motor-evoked potentials by TMS (Mhalla et al., 2010), were also detectable globally through GMFP, and in the stimulated area with LMFP, and peak-to-peak amplitudes using TMS-EEG. Secondly, we investigated whether changes in alpha-band oscillatory dynamics, particularly ERSP and ITC, previously observed in experimental pain models (De Martino et al., 2024), were also reduced in individuals with FM. Finally, it was examined whether individuals with FM exhibit reduced natural frequency and measures of cortical complexity, potentially reflecting the disrupted spatiotemporal organization of cortical responses. Additionally, correlations between the potentially observed changes in individuals with FM and their main clinical characteristics of FM were explored.

## 2. METHODS

### 2.1. Participants

This study was conducted at the Center for Neuroplasticity and Pain (CNAP) at Aalborg University from April-November 2024. The protocol was approved by the Ethics Review Board (N-20230076/N-2023007) and complied with the declaration of Helsinki.

Individuals diagnosed with FM based on the ACR 2016 criteria (Wolfe et al., 2016) and pain-free controls were included. Inclusion criteria for individuals with FM were age 18-80 years old, speak and understand English or Danish, and have an average pain intensity above 3 for most days of the week on a 0-10 pain intensity numerical rating scale (where 0 is no pain and 10 is the worst possible pain) (Cleeland, 2009). Inclusion criteria for pain-free controls were age between 18-80 years old and had to speak and understand English or Danish. Exclusion criteria for both groups were pregnancy and/or breastfeeding, current diagnoses of major depression or drug abuse (alcohol or opioids), lack of ability to cooperate and trouble answering the questionnaires (due to language or cognitive problems), participating in another trial in the month before inclusion and any contraindications to TMS (e.g., brain metal implants, uncontrolled epilepsy) (Rossi et al., 2021). Additional exclusion criteria for controls were the presence of chronic pain (pain most days within the last three months), known neurologic and musculoskeletal diseases, psychiatric diagnoses, use of analgesic substances, episodes of acute pain, or abnormal sleep within the previous 48 hours before assessment.

A sample size calculation was performed using GPower (G*Power 3.1.9.7, Düsseldorf, 2020) for an independent samples t-test to compare individuals with FM and controls. Based on a previous study investigating intracortical inhibition by paired pulses of TMS to the left M1 (Mhalla et al., 2010), individuals with FM demonstrated reduced intracortical inhibition (mean ± SD = 60 ± 20%), while the control group showed 80 ± 20% inhibition after investigating motor evoked potentials by TMS. Assuming a significance level (α) of 0.05 and a power of 0.80, the effect size was 1.0, resulting in a required sample size of 14 participants per group in this study. To account for possible technical challenges, about 25% of additional participants were recruited for each group.

### 2.2. Study Design

This report presents results from the baseline assessment of individuals with FM included in an ongoing clinical trial on the use of repetitive TMS for chronic pain (Clinicaltrials.gov: NCT06395649). All participants participated in a single experimental session lasting approximately three hours. First, FM and control participants filled out questionnaires and subsequently underwent EEG recordings under single pulse TMS delivery to the hand area of the left M1.

### 2.3. Demographics and Clinical assessment

All participants reported demographic data (age, sex, height, and weight). Individuals with FM also reported pain duration and current medication intake. The type and dosing of medication for each participant were calculated based on the Medication Quantification Scale in which medications are scored based on their propensity to cause mental confusion, sleepiness, or central side effects (Harden et al., 2005). Individuals with FM completed the Short Form Brief Pain Inventory questionnaire (BPI) (Cleeland & Ryan, 1994; Cleeland, 2009). The BPI measures pain severity (current, worst, least, and average pain within the previous 24 hours, each ranging from 0 to 10; no pain to worst pain imaginable) and pain interference with daily function (general activity, mood, ability to walk, normal work, relationship with other people, sleep and enjoyment of life). Each of the seven pain interference items are scored from 0 to 10 (does not interfere to completely interfere). BPI also provides a list of current ongoing pain therapies and the average pain relief patients had under their use (0-100% pain relief) (Cleeland, 2009). Mood and anxiety were assessed with the 14-item Hospital Anxiety and Depression Scale (HADS) which includes 7 items for anxiety and 7 for depression. Higher scores indicate more severe symptoms (Bjelland et al., 2002; Zigmond & Snaith, 1983). The EuroQol-5 Domain (EQ-5D) was used to assess quality of life involving domains of mobility, personal care, usual activities, pain, and anxiety/depression, ranging from 1-5, with a total score of 5 being the lowest score. A lower score indicates a higher quality of life (Herdman et al., 2011).

### 2.4. TMS-EEG Procedure

EEG was measured using an amplifier (g.HIamp EEG amplifier, g.tec medical engineering GmbH) with a passive electrode cap containing 64 electrodes (Easycap) placed according to the 10-5 system, with the Cz electrode on the vertex. The ground electrode was placed on the right zygoma, the online reference was on the right mastoid process, and two electrodes were placed on the lateral side of the eyes to record eye movement. Electrode impedance was kept under 5 kΩ (Groppa et al., 2012). Raw signals were amplified and sampled at a rate of 4800 Hz. To avoid the electrodes from drying out, to keep them in contact with the scalp, and to prevent somatosensory artifacts produced by the coil touching the electrodes, two net caps was placed on top of the EEG-cap and afterwards wrapped with plastic (Hernandez-Pavon et al., 2023). To prevent auditory EEG responses occurring due to the click-sounds from the TMS coil during stimulation, participants wore earplugs with noise-cancelation connected to the TMS-click sound masking toolbox (TAAC) (Russo et al., 2022).

Participants were slightly reclined in a comfortable chair with the head rested back and instructed to keep their eyes open and relax as much as possible. TMS-pulses were applied (MagPro R30, MagVenture A/S, Farum, Denmark) with a figure-of-eight coil (Cool-B35) to the scalp projection of left M1. The targeted hotspot was defined by the coil position that evoked a maximal visual motor response for a given stimulation intensity (Rossini et al., 2015). Given the hotspot, the resting motor threshold (RMT) was estimated by lowering the stimulation intensity until a contraction of the right hand was no longer observed (De Martino, Nascimento Couto et al., 2025). Real-time TMS evoked EEG potentials (TEPs) recordings were used to adjust the position and orientation of the coil, as well as the TMS intensity, to minimize artifacts, and ensure a sufficient amplitude as measured on the peak-to-peak over the electrodes below the coil (C1, C3, CP1 and CP3) (Casarotto et al., 2022). Stimulation intensity was initially set to 90 % of the RMT. To monitor the coil position during pulse delivery, a navigate brain stimulation software (InVesalius Navigator, 3.1) was used with an optical (infrared) motion capture camera (Polaris Vicra, NDI, Ontario, Canada) (Amorim et al., 2015). An average of 20-60 pulses were given before the actual recording to ensure signal quality, and approximately 200 pulses were delivered for the actual experimental recording with interstimulus intervals randomly jittered between 2000 and 2500 ms (Casarotto et al., 2010).

### 2.5. Pre-processing of TMS-EEG

Artifacts produced by the TMS-pulse 0 to 15 ms after each stimulation were removed and replaced by the temporally reversed pre-pulse signal (-15 to 0 ms). A high-pass filter (1 Hz) was applied to remove signal drift. TEPs were segmented from –800 to 800 ms around the TMS pulse onset. All TEPs from all channels were visually inspected and artefactual epochs (e.g., coil movement, eye blinks, and muscle activity) were removed (De Martino, Casali et al., 2025). Independent component analysis (ICA) was used to remove residual artifacts (e.g., eye movements, eye blinks, muscle activity, 50 Hz, decay artifacts) (Delorme & Makeig, 2004). Then, a bandpass filter (1–80 Hz, Butterworth, 3^rd^ order) was applied and EEG data were down sampled from 4800 Hz to 1200 Hz. Channels removed during visual inspection were interpolated, the reference was set to the average of all channels, and the data underwent baseline correction, as previous described (De Martino, Nascimento Couto et al., 2025). TEPs with signal-to-noise-ratio below 1.4 were excluded from the data analysis (Casali et al., 2013). For all analyses the grand average TEP per channel were calculated, expect for the ERSP.

### 2.6. TMS Evoked Potentials Outcomes

#### 2.6.1. Cortical excitability

To assess global cortical excitability to TMS, GMFP was calculated as the mean square root power across all EEG channels of the 20-300 ms post-TMS pulse (De Martino et al., 2023). Local cortical excitability was assessed using the LMFP of the 20-300 ms post-TMS pulse, computed from electrodes cluster positioned under the stimulation site (C1, C3, CP1, and CP3). LMFP and GMFP results are presented in mV/sec. Additionally, to assess the early responses in the stimulated area, the peak-to-peak amplitude of TEPs was quantified by measuring the difference between the first positive peak (P30) and the first negative peak (N100) at C1, C3, CP1, and CP3 electrodes. These peak-to-peak amplitudes were then averaged across the local electrode cluster. Latency was extracted by subtracting the timepoint P30 from N100.

#### 2.6.2. Oscillatory activity

Time-frequency representation of the data was computed using Morlet wavelets across a frequency range of 8–45 Hz (3.5 cycles, 1 Hz resolution) (Cohen, 2014; Delorme & Makeig, 2004). Statistical significance of evoked oscillatory activity was assessed with a two-tailed bootstrap statistic. Null distributions for both Event-Related Spectral Perturbation (ERSP) and Inter-Trial Coherence (ITC) were generated by creating 500 surrogate datasets from shuffled pre-stimulus activity.

The ERSP value was calculated by averaging the time-frequency maps over a region of interest directly under the TMS coil (C1, C3, CP1, CP3). From this analysis, we identified the natural frequency by finding the peak power in the spectrum within the 20−300 ms window after the TMS pulse (Rosanova et al., 2009).

Next, guided by previous reports on the link between oscillatory connectivity and pain sensitivity (De Martino et al., 2024), we performed a more targeted examination of oscillatory activity. We analyzed the same 20−300 ms post-stimulation interval but focused on specific electrode clusters and frequency bands identified in that prior research. ERSP was examined in a medio-frontal cluster (FCz, Cz, FC1, FC2, C1, C2) and ITC in alpha-band parietal occipital (PO) clusters: left (left PO: PO7, PO3, PO1), midline (middle POz, Oz, PO3, PO4), and right (right PO: PO8, PO4, O2).

#### 2.6.3 Perturbational complexity

The PCIst was derived from TMS-EEG evoked potential through a series of steps. First, TEPs were decomposed into *N_c_* principal components using singular value decomposition. Then, amplitude distances were calculated for each component between baseline and post-pulse response samples, forming respective distance matrices. These matrices were then thresholded at multiple scales, defining different transition levels (i.e., different state transitions between different levels of sensitivity). At each scale, transition matrices were computed for both baseline and post pulse response, allowing for the estimation of state transitions (*NST*) in each condition. The complexity of each component was determined by the maximum weighted difference between response and baseline transitions (Δ*NST_n_*) and the final perturbational complexity index measure was obtained by summing Δ*NST_n_* across all principal components (Comolatti et al., 2019).

### 2.7. Statistics

Descriptive statistics were performed in STATA (StataCorp. 2023. *Stata Statistical Software: Release 18*. College Station, TX: StataCorp LLC). All data were assessed for normality by visual inspection of histograms and the Shapiro-Wilk test and presented as median with interquartile range [25^th^ and 75^th^ percentiles], if not otherwise stated. TMS-EEG data were compared between the two groups. If the data were normally distributed and followed the assumptions for a parametric test, the Student’s unpaired t-test was used and otherwise the non-parametric Mann-Whitney U test was used. Parameters that were significantly different between the two groups were assessed for correlation with clinical characteristics by Spearman’s Rank-Order Correlation within the individuals with. Due to the exploratory nature of this study and its hypothesis generating rather than confirmatory aim, corrections for multiple comparisons were not undertaken (Bender & Lange, 2001; Streiner, 2015).

## 3. Results

Twenty individuals with FM and 16 pain-free controls were included in this study. The data from two participants with FM and one control were excluded from the analysis as the TMS-EEG signal-to-noise-ratio was <1.4. Individuals with FM reported pain intensities of 6.4 [5.8-6.8], pain interference of 6.1 [4.4-7.0], and a Medication Quantification Scale score of 8.4 [3.8-13.5] with 4/18 not taking any medication (see supplementary material for specific drugs). Other pain characteristics and sociodemographic details are displayed in Table 1.

**Table 1.** Demographic data of all participants and clinical characteristics of individuals with fibromyalgia. All data, except sex, are expressed as median with interquartile range [25th and 75^th^percentiles] and the minimum and maximum value (min-max). * = p < 0.05.

| Demographic data | Fibromyalgia | Pain-free controls | P-value |
| --- | --- | --- | --- |
| **General** |  |  |  |
| Sex (women) | 18/18 | 10/15 | - |
| Age (years) | 52.5 [42.0-57.0] (25.0-63.0) | 51.0 [39.0-59.0] (24.0-71.0) | 0.871 |
| Height (cm) | 168.5 [164.0-173.0] (162.0-178.0) | 168.0 [162.0-175.0] (160.0-206.0) | 0.651 |
| Weight (kg) | 90.0 [70.0-92.0] (58.0-105.0) | 77.0 [70.0-80.0] (52.0-104.0) | 0.124 |
| Pain duration (years) | 13.0 [10.0-17.0] (4.0-30.0) | - | - |
| Number of drugs for pain | 2.0 [1.0-3.1] (0.0-8.0) | - | - |
| Medication Quantification Scale | 8.4 [3.8-13.5] (0.0-34.4) | - | - |
| **Pain intensity - BPI** |  |  |  |
| BPI Pain Intensity (NRS) Total Score | 6.4 [5.8-6.8] (3.5-7.3) | - | - |
| Worst pain | 8.0 [8.0-9.0] (6.0-9.0) | - | - |
| Least pain | 4.0 [3.0-5.0] (2.0-9.0) | - | - |
| Average pain | 6.0 [6.0-7.0] (4.0-8.0) | - | - |
| Pain right now | 6.0 [5.0-7.0] (2.0-8.0) | - | - |
| Relief of pain due to therapy (%) | 40.0 [30.0-50.0] (0.0-100.0) | - | - |
| **Pain interference - BPI** |  |  |  |
| BPI Pain Interference (NRS) Total Score | 6.1 [4.4-7.0] (3.6-8.3) | - | - |
| General activity | 6.5 [6.0-7.0] (2.0-8.0) | - | - |
| Mood | 6.0 [4.0-7.0] (2.0-8.0) | - | - |
| Walking ability | 6.0 [4.0-7.0] (0.0-9.0) | - | - |
| Normal work (including housework) | 7.0 [5.0-8.0] (2.0-10.0) | - | - |
| Relationship with other people | 6.0 [3.0-7.0] (2.0-10.0) | - | - |
| Sleep | 7.5 [4.0-9.0] (0.0-10.0) | - | - |
| Taste for life | 7.0 [4.0-8.0] (2.0-9.0) | - | - |
| **Quality of life – EQ-5D** |  |  |  |
| Total Score | 11.0 [10.0-14.0] (8.0-15.0) | 5.0 [5.0-5.0] (5.0-7.0) | 0.000* |
| Mobility | 2.0 [2.0-3.0] (1.0-4.0) | 1.0 [1.0-1.0] (1.0-1.0) | 0.000* |
| Selfcare | 1.0 [2.0-2.0] (1.0-3.0) | 1.0 [1.0-1.0] (1.0-1.0) | 0.008 |
| Usual activities | 3.0 [3.0-4.0] (1.0-4.0) | 1.0 [1.0-1.0] (1.0-2.0) | 0.000* |
| Pain/discomfort | 3.0 [3.0-4.0] (3.0-4.0) | 1.0 [1.0-1.0] (1.0-2.0) | 0.000* |
| Anxiety/depression | 1.0 [1.0-2.0] (1.0-3.0) | 1.0 [1.0-1.0] (1.0-1.0) | 0.006 |
| **HADS** |  |  |  |
| Anxiety Total Score | 7.0 [6.0-8.0] (0.0-17) | 3.0 [1.0-4.1] (0.0-9.0) | 0.003* |
| Depression Total Score | 5.0 [4.0-8.0] (0.0-10.0) | 0.5 [0.0-1.0] (0.0-4.0) | 0.000* |

Abbreviations: The Short Form of Brief Pain Inventory questionnaire (BPI), EuroQol-5 Domain questionnaire (EQ-5D), Hospital Anxiety and Depression questionnaire (HADS), Numerical Rating Scale (NRS).

### 3.1. Cortical excitability

RMT for the FM group was 53.5 [49.0-57.0] % and 50.0 [42.0-55.0] % for the pain-free controls which was not significantly different (p=0.118). Individuals with FM showed no significant difference in the GMFP (Table 2): FM 0.37 [0.25-0.41], controls 0.29 [0.22-0.42]; p=0.343 or in the LMFP: FM 0.38 [0.34-0.53] when comparing with controls 0.39 [0.25-0.44] (p=0.662). Individuals with FM had a TEP peak-to-peak of 7.00 [4.83-7.77] µV and a peak-to-peak-latency of 36.88 [25.75-49.00] ms. These measurements were not significantly different from controls, who had a peak-to-peak amplitude of 6.51 [4.38-10.17] µV (p=0.957) and latency of 32.75 [28.00-40.50] ms (p=0.662).

**Table 2.** Transcranial Magnetic Stimulation (TMS) evoked potentials outcomes. All data are expressed as median with interquartile range [25th and 75th percentiles] and the minimum and maximum value (min-max). * = p < 0.05. Peak-to-peak and natural frequency were measured at electrodes below the coil (C1, C3, CP1, CP3).

| *TMS Evoked Potentials Outcomes* | *Fibromyalgia* | *Pain-free controls* | *P-value* |
| --- | --- | --- | --- |
| Peak-to-peak amplitude | 7.00 [4.83-7.77]<br>(2.92-12.12) | 6.51 [4.38-10.17]<br>(2.45-21.01) | 0.942 |
| Peak-to-peak latency | 36.88 [25.75-49.00]<br>(20.00-75.50) | 32.75 [28.00-40.50]<br>(13.50-54.50) | 0.651 |
| GMFP | 0.37 [0.25-0.41],<br>(0.17-0.52) | 0.29 [0.22-0.42]<br>(0.18-0.59) | 0.329 |
| ERSP MF | 0.04 [-0.02-0.10]<br>(-0.06-0.25) | 0.10 [0.00-0.13]<br>(-0.01-0.28) | 0.233 |
| Left PO ITC | 0.15 [0.11-0.13]<br>(0.06-0.38) | 0.24 [0.14-0.27]<br>(0.07-0.39) | 0.055 |
| Middle PO ITC | 0.13 [0.10-0.20]<br>(0.06-0.27) | 0.22 [0.15-0.23]<br>(0.07-0.32) | 0.043* |
| Right PO ITC | 0.10 [0.08-0.14]<br>(0.05-0.24) | 0.18 [0.10-0.21]<br>(0.05-0.28) | 0.039* |
| Natural frequency | 11.00 [9.00-13.00]<br>(8.00-17.00) | 11.00 [10.00-13.00]<br>(8.00-23.00) | 0.688 |
| Perturbational complexity index | 31.71 [24.75-34.74]<br>(19.66-53.15) | 41.21 [33.93-47.54]<br>(16.31-56.47) | 0.010* |

Abbreviations: GMFP: Global mean field power, ERSP: Event-related spectral perturbation, MF: Medio-frontal cluster, PO: Parietal occipital cluster, ITC: Intertrial coherence.

### 3.2. Oscillatory activity

Alpha-band ITC was significantly lower in individuals with FM than controls on the middle and right PO (Figure 1A and 1B, respectively): middle PO, FM= 0.13 [0.10-0.18], controls= 0.22 [0.15-0.23]; p=0.044, right PO, FM=0.10 [0.08-0.14], controls= 0.18 [0.10-0.25]; p= 0.039. Alpha-band ITC on the left PO was lower in individuals with FM compared to controls but not significantly different: FM=0.15 [0.11-0.18], controls 0.24 [0.14-0.27]; p=0.057 (Figure 1.C). No significant difference was found in the natural frequency of M1 between individuals with FM 11.00 [9.00-13.00] Hz and controls (11.00 [10.00-13.00] Hz), (p=0.696), as well as in the alpha-band ERSP in the medio-frontal cluster (FM= 0.04 [-0.02-0.10], controls=0.10 [0.00-0.13]), (p=0.244).

**Figure 1.**
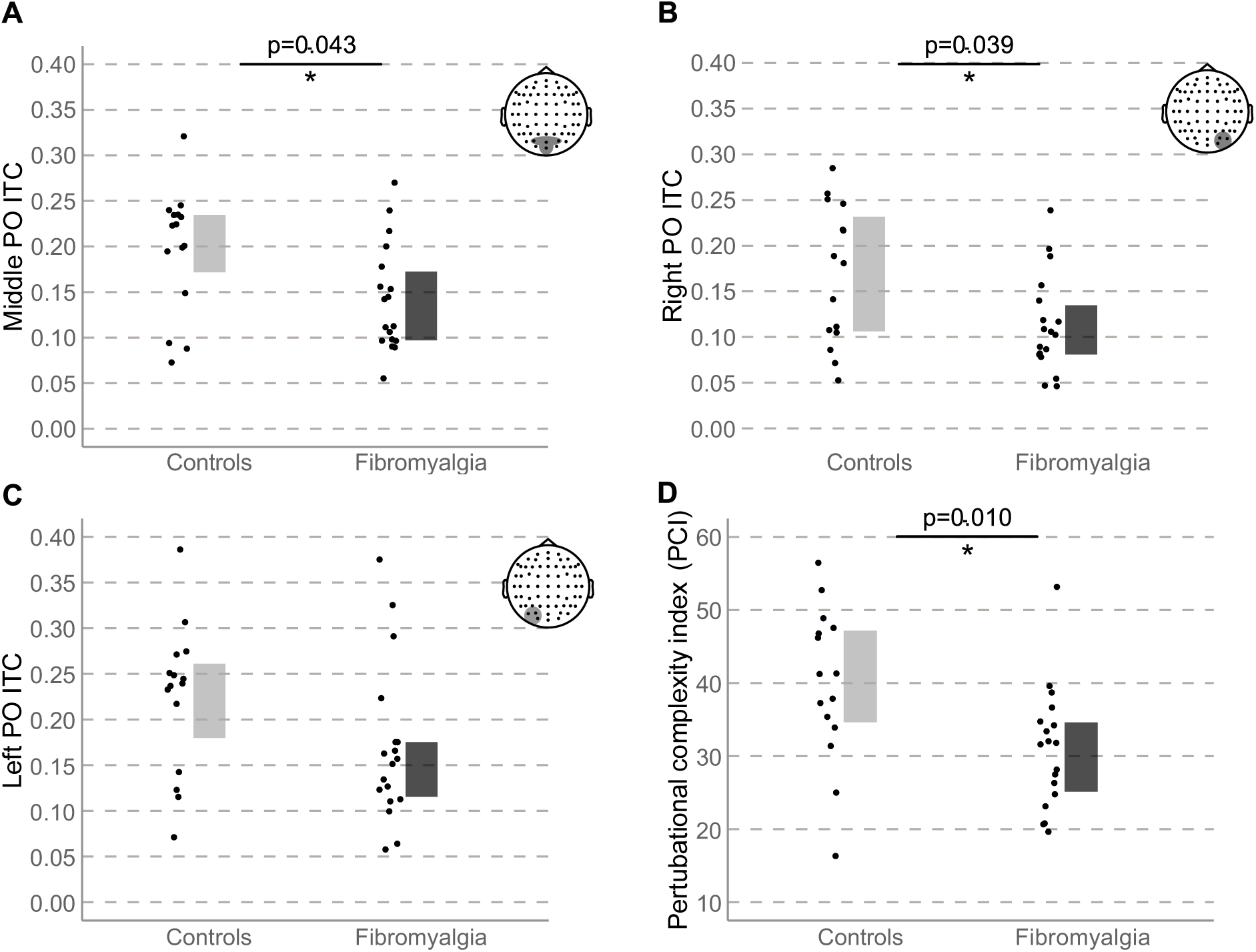
Comparison of main results between individuals with fibromyalgia and pain-free controls showing. A) middle parietal occipital (PO) intertrial coherence (ITC), B) right PO ITC, C) left PO ITC and D) Perturbational complexity index (PCI). All figures show individual data points, boxplot with median and interquartiles and the distribution of the data. On A, B, and C the topoplot on the superior right corner illustrates the cluster used for each comparison. * = p < 0.05.

### 3.3. Perturbational complexity

The PCIst was significantly lower in individuals with FM (31.71 [24.75-34.74]) than in controls (41.21 [33.93-47.54]); p=0.009 (Figure 1.D).

### 3.4. Correlation analysis

In the FM group, alpha-band middle-PO ITC negatively correlated with percentage of pain relief with the current therapy (rho= -0.552, p=0.019; Figure 2.A). The PCIst showed a negative correlation with the interference of pain (rho=-0.486, p=0.042; Figure 2.B) and a negative correlation with percentage of pain relief with the current therapy (rho= -0.478, p=0.046; Figure 2.C). All other relevant correlations were non-significant.

**Figure 2.**
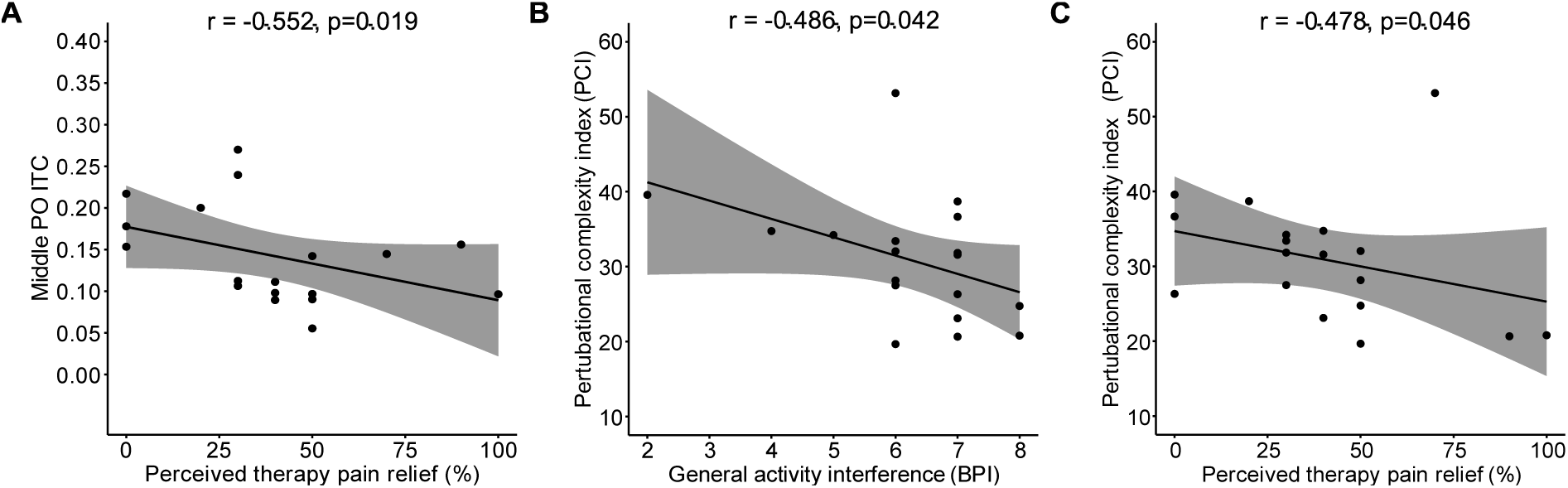
Significant correlations between Transcranial Magnetic Stimulation (TMS) evoked potentials and clinical outcomes for the fibromyalgia group. A) middle parietal occipital (PO) intertrial coherence (ITC) and perceived therapy pain relief (measured on the Short Form Brief Pain Inventory questionnaire (BPI)), B) Perturbational complexity index (PCI) and pain interference with general activity level (BPI, interference) and C) PCI and perceived therapy pain relief. All correlations were significant (p<0.05). Only outcomes where there was a significant difference between the pain-free and fibromyalgia groups are shown.

## 4. DISCUSSION

Individuals with FM had decreased alpha-ITC in parietal occipital areas, and a lower capacity to generate complex oscillatory dynamics following M1 TMS perturbation. Alpha-ITC and PCIst correlated with pain interference and percentage of pain relief with the current therapy of FM and were unlikely to be related to medication use or dosage. Cortical excitability metrics did not significantly differ between individuals with FM compared to healthy controls.

### 4.1. Cortical Excitability

Two measurements of local cortical excitability were assessed in the current study: the LMFP and the peak-to-peak amplitude. LMFP reflects the local cortical response of 4 EEG electrodes near M1 over a time window (20–300 ms), primarily indicating local cortical excitability (Komssi & Kähkönen, 2006). In this measure, no significant differences were observed in individuals with FM. These findings contrast with those of previous experimental pain studies where acute pain has been reported to influence excitability measures in healthy people (De Martino et al., 2023). In individuals with FM, a reduction in corticospinal excitability, as assessed by MEP and bilateral decreases in intracortical facilitation and short intracortical inhibition have been reported (Mhalla et al., 2010). One possible explanation for these differing findings is that spinal and peripheral mechanisms may influence some of the MEP-based measures (Di Lazzaro & Ziemann, 2013). In contrast to these studies, we found no differences in RMT between pain-free individuals and individuals with fibromyalgia. Additionally, TMS-EEG used in the current study was below motor threshold, allowing for a more direct assessment of cortico-cortical and corticothalamic circuits.

A TMS-pulse leads to an oscillatory evoked response containing several peaks (Komssi et al., 2004). It has been suggested that earlier peaks (N15-P30) are linked to cortical excitatory activity, while later peaks (N45-N100) refer to cortical inhibition. By measuring the peak-to-peak amplitude at certain electrodes, the response of this specific area to the TEP can be quantified (Hill et al., 2016). During experimental acute pain, a reduction in the first peak-to-peak responses were observed compared to non-noxious warm stimuli when stimulating M1 and dorsolateral prefrontal cortex (DLPFC) by TMS in the respective stimulation areas (De Martino et al., 2023). Acute heat pain has also been reported to increase frontal TEP components, such as the N45 peak. This increase in amplitude was correlated with higher pain ratings and associated with changes in cortical excitability (Chowdhury et al., 2023). The N45 appears approximately 45 milliseconds after the TMS pulse and is thought to reflect GABAergic activity and inhibitory control (Premoli et al., 2014). The absence of peak amplitude changes in the current study supports the idea that acute experimental pain induces changes in cortical excitability that are different from chronic pain states, which corresponds to the differing mechanistic and phenomenological experiences between these two distinct experiences.

GMFP captures the overall TMS-evoked response by averaging activity across all electrodes over a broad time window (20–300 ms in this study), thereby reflecting global cortical excitability and incorporating aspects of functional connectivity (Tremblay et al., 2019). In pain-free individuals under experimental pain, nociceptive heat was shown to reduce GMFP compared with baseline and non-noxious warm stimuli within the same time interval of the present study (De Martino et al., 2023), suggesting that acute pain may affect global cortical excitability. In the current study, no significant reduction in GMFP was observed in individuals with FM compared to healthy controls. However, given the cross-sectional nature of this study, it remains unclear whether GMFP changes occur during the natural history of the condition or if the widespread nature of FM pain gives different manifestations compared to localized experimental pain.

### 4.2 Oscillatory Dynamics

ITC is a phase-based functional connectivity measurement. It measures how consistent brain oscillations are across repetitive trials under a certain time and frequency interval (Makeig et al., 2004). Acute experimental heat pain has been shown to lead to reductions in the normal expected ITC within this same regiońs natural oscillatory behavior, the alpha-band (De Martino et al., 2024). The reductions in ITC correlated during acute pain with the sensitivity to pain, indicated that people with lower pain thresholds (i.e., more sensitive to pain) showed more marked reductions in phase synchronization during acute pain, while those who had higher pain thresholds (i.e., less sensitive to pain) exhibited less interference in alpha-band ITC (De Martino et al., 2024). In the present study, such decreases in ITC were found to occur in individuals with FM within the same area and frequency band, when compared to pain-free controls. Additionally, phase coherence changes negatively correlated with the subjective rating of pain relief under the current therapy, so that those experiencing more pain relief with treatment had more reductions in ITC. It is important to note that all patients were under at least moderate pain at the time of assessment. Perceived pain relief was evaluated based on subjective reports of changes in the participants’ overall pain state after starting their current therapy. These findings suggest that reductions in normal posterior alpha phase-based coherence may be linked to how individuals appraise their typical pain sensitivity, or trait. Thus, stoic people had less interference (reductions) in their pain-free ITC results when they experienced acute pain (De Martino et al., 2024), and ITC reductions more marked in patients who consider having had more pain relief so far. In this view, reductions in ITC would correlate with an individual’s appraisal of their higher pain sensitivity or “functional reserve” to integrate nociception. It has been suggested that alpha oscillatory activity is influenced by thalamocortical GABAergic projections (Lim et al., 2016; Sarnthein et al., 2006). GABA-A receptors are believed to be key players in the generation of oscillatory activity in lamina V pyramidal neurons in the brain, and disruption of their activity may lead to abnormal oscillatory activity and abnormal connectivity (Barbero-Castillo et al., 2021). Interestingly, both healthy participants experiencing experimentally induced acute pain and individuals with FM have been reported to exhibit defective GABA-A dependent intracortical inhibition (Chowdhury et al., 2023; Mhalla et al., 2010). Intracortical inhibition correlated with symptoms in FM individuals (Mhalla et al., 2010), and was partially restored by therapy (Mhalla et al., 2011).

Cortical oscillatory dynamics can also be measured by changes in power of evoked responses (Rosanova et al., 2009). The ERSP enables identification of power changes over time and across frequencies after an event (Makeig et al., 2004). In the present study, no significant alpha-band ERSP differences were observed in the medio-frontal cluster, again, contrasting with findings from experimental pain studies that reported reductions in ERSP (De Martino et al., 2024). This further highlights differences between acute pain and clinical pain, also reported in cortico-spinal excitability measures based on MEP-based excitability measures.

No significant between-group differences were found in the natural frequency, which refers to the dominant frequency of cortical responses following the TMS-evoked potential (Rosanova et al., 2009). The natural frequency in individuals with schizophrenia (Donati et al., 2025; Ferrarelli et al., 2008) major depression, and bipolar disorder is known to be reduced compared to pain-free controls (Canali et al., 2015), suggesting disruption of the normal dynamics of oscillatory cortical activity, altering the normal dominant frequency band expected in discrete brain cortical areas in these disorders. Even though depressive symptoms can be a part of FM, and it has been suggested that both disorders have similar traits (Yepez et al., 2022), results from the present study suggest that FM effects on brain dynamics are different from those of major depressive disorder.

### 4.3. Perturbation Complexity Index

The PCIst is a measure of the balance between integration and differentiation of thalamocortical, reflecting the degree of complexity of responses following a perturbating pulse of TMS. It has been studied as an objective tool to assess consciousness in individual patients (Casali et al., 2013; Comolatti et al., 2019). Scores of the PCIst have been reported to be higher in healthy awake people when compared to values under non-rapid eye movement sleep and even lower for patients under anesthesia (Casali et al., 2013; Casarotto et al., 2016). Similarly, in patients recovering from coma and reaching an intermediate level of consciousness, the PCIst is significantly influenced by the patient’s level of consciousness, where higher scores predicted higher chances of patients developing higher consciousness levels on follow-up, some also gaining content of consciousness (Casarotto et al., 2016). In the present study, lower values of the PCIst based on state transitions were found in individuals with FM compared to pain-free controls. The PCIst is expected to be low when multiple interacting brain regions respond to the TMS perturbation in a highly stereotyped manner, leading to a loss of differentiation, or due to reduced interaction among regional cortical areas (Comolatti et al., 2019). It is possible that in FM low PCIst values reflected a reduction in connectivity within M1, or perhaps between other regions, thus affecting the result of the index. Given the normal LMFP and peak-to-peak amplitudes, the latter possibility seems more likely. Importantly, when assessing patients in coma or with reduced levels of consciousness, healthy conscious individuals exhibited higher complexity, with PCIst values spanning a wide range (∼21–80, mean 47.89 ± 12.65), whereas loss of consciousness was consistently associated with low PCIst values (below ∼21) (Comolatti et al., 2019). To date, there is no evidence that PCIst values above this cut-off, yet on the lower end, carry any biological relevance. To date, there is still not enough data suggesting that lower PCIst values that are still above these cut-off values would have any biological relevance. The assessment of other complexity measures such as entropy-based ones, or approaches assessing fractal dimension and near chaotic behavior would better help clarify the relevance of reduced PCIst results in FM. These next steps would allow to confirm whether individuals with FM have lower than average perturbational complexity indices, and if so, whether such changes are reflecting the effects of psychotropic medication, FM-specific decreased consciousness levels in a delirium-like pattern (Bai et al., 2023) such as “fibro-fog”, or broader decreases in global connectivity.

### 4.4. Limitations

Firstly, over 75 % of the individuals with FM were taking some medication. Although medication does not seem to correlate with ITC and PCIst in the present study and that alterations in MEPs do not seem to be linked to medication use in FM (Mhalla et al., 2010), it cannot be excluded that medication affect the TEPs. Secondly, future studies could benefit from an assessment of cognitive performance in individuals with FM, due to the large impact of cognition in the patient group (Bell et al., 2018). It cannot be excluded that cognition could have an impact on TMS evoked measurements in individuals with FM, as assessments of individuals with major depression have found a negative association between P30 amplitude and cognitive dysfunction (Li et al., 2024).

Concerning sex differences, it is known that men with FM are more likely to present with certain medical comorbidities, such as obstructive sleep apnoea and alcohol use disorders, as well as a higher prevalence of post-traumatic stress disorder (Bannon et al., 2025; Drusko et al., 2023). Also, population-based data suggest that the association between FM and major depressive disorder and generalized anxiety disorder is stronger in men, indicating a potentially gender-specific pattern of comorbidity (Thomas et al., 2024). Some studies also suggest that men may have a higher prevalence of substance use disorders and may employ different pain coping strategies compared to women (Bannon et al., 2025; Drusko et al., 2023). Here we decided to study exclusively women with FM, and compared them to a control group, as large normative data indicate no significant gender-specific differences in cortical excitability measures, such as MEPs (Cueva et al., 2016). Even if higher variability in the control group can reduce statistical power, we could still detect differences between groups in a relatively small sample of patients.

Lastly, as this was an exploratory cross-sectional study, it remains unclear whether the observed lower ITC and PCIst in individuals with FM compared to controls preceded the onset of symptoms or emerged because of FM symptoms, and whether these alterations may evolve over time (Wang & Cheng, 2020). All the reticence related to absence of correction for multiple comparisons need to be made and our results need to be interpreted as an initial exploration of advanced neurophysiology exploration of people with primary pains by using TMS-EEG.

### 4.5 Conclusions

In this study, no significant differences were found in cortical excitability measures between individuals with FM compared to pain-free controls. However, individuals with FM showed a reduction in alpha-band oscillatory coherence and broad decreases in global connectivity. These findings, if replicated, could help better understand the brain dynamics in FM.

## Acknowledgements

The Center for Neuroplasticity and Pain (CNAP) is supported by the Danish National Research Foundation (DNRF121). Relevant to this study, Thomas Graven-Nielsen has received funding from the Lundbeck Foundation (R441-2023-232). This research has received funding from the European Research Council (ERC) under the European Union’s Consolidator Grant: PersoNINpain (DCA, EDM, BANC, AJ, MMB) (grant agreement No. 101040123) and Novo Nordisk Start Package for Faculty Enrollment 223124 (DCA, AJ, EDM, SIM).

