## Supplementary material 1 for "REDUCED ALPHA-BAND PHASE COHERENCE AND CORTICAL COMPLEXITY IN FIBROMYALGIA: A TMS-EEG EXPLORATORY STUDY"

**Supplementary material 1:** Number of individuals with fibromyalgia taking drugs and name of the drug.

| **Drug:** | **Number of individuals:** |
|---|---|
| Amitriptylin | 1 |
| Baclofen | 1 |
| Citalopram | 2 |
| Duloxetine | 2 |
| Gabapentin | 2 |
| Hydroxyzine | 1 |
| Ibuprofen | 6 |
| Lyrica (same as pregabalin) | 1 |
| Methylphenidat | 1 |
| Mirtazapin (antidepressive) | 1 |
| Morfin | 1 |
| Paracetamol | 10 |
| Paroxetin | 1 |
| Pregabalin | 2 |
| Quetiapin | 1 |
| Sertraline (anti | 1 |
| Sifrol | 2 |
| Tramadol | 5 |
